# Summer temperature can predict the distribution of wild yeast populations

**DOI:** 10.1101/027433

**Authors:** Heather A. Robinson, Ana Pinharanda, Douda Bensasson

**Author notes:** Author for correspondence: Douda Bensasson Faculty of Life Sciences University of Manchester Michael Smith Building Oxford Road Manchester, M13 9PT United Kingdom. Present address: Department of Zoology, University of Cambridge (Ana Pinharanda).

## Abstract

The wine yeast, *Saccharomyces cerevisiae,* is the best understood microbial eukaryote at the molecular and cellular level, yet its natural geographic distribution is unknown. Here we report the results of a field survey for *S. cerevisiae, S. paradoxus* and other budding yeast on oak trees in Europe. We show that yeast species differ in their geographic distributions, and investigated which ecological variables can predict the isolation rate of *S. paradoxus*, the most abundant species. We find a positive association between trunk girth and *S. paradoxus* abundance suggesting that older trees harbour more yeast. *S. paradoxus* isolation frequency is also associated with summer temperature, showing highest isolation rates at intermediate temperatures. Using our statistical model, we estimated a range of summer temperatures at which we expect high *S. paradoxus* isolation rates, and show that the geographic distribution predicted by this optimum temperature range is consistent with the worldwide distribution of sites where *S. paradoxus* has been isolated. Using laboratory estimates of optimal growth temperatures for *S. cerevisiae* relative to *S. paradoxus*, we also estimated an optimum range of summer temperatures for *S. cerevisiae*. The geographical distribution of these optimum temperatures are consistent with the locations where wild *S. cerevisiae* have been reported, and can explain why only human-associated *S. cerevisiae* strains are isolated at northernmost latitudes. Our results provide a starting point for targeted isolation of *S. cerevisiae* from natural habitats, which could lead to a better understanding of climate associations and natural history in this important model microbe.

## Introduction

The wine yeast, *Saccharomyces cerevisiae* is of considerable importance to humans for agriculture, industry, and basic research, but little is known about its ecology (Goddard and Greig, 2015; Liti, 2015). Wild populations of *S. cerevisiae* have been isolated from oak and other tree species in North America, Europe and Asia (Sniegowski *et al.*, 2002; Sampaio and Gonçalves, 2008; Diezmann and Dietrich, 2009; Wang *et al.*, 2012; Hyma and Fay, 2013), and are genetically distinct from those associated with human activity (Fay and Benavides, 2005; Cromie *et al.*, 2013; Almeida *et al.*, 2015). These woodland habitats and the populations they contain therefore represent a good target for revealing the ecology of *S. cerevisiae,* and the full extent of phenotypic and genetic diversity within the species. A fundamental challenge, however, is that the natural geographic distribution of *S. cerevisiae* is unknown. Indeed, geographic distributions are described for only few individual, free-living microbial species (Taylor *et al.*, 2006; Green and Bohannan, 2006; Martiny *et al.*, 2006). In Portugal and parts of the USA, *S. cerevisiae* is sympatric with *S. paradoxus* (Sniegowski *et al.*, 2002; Sampaio and Gonçalves, 2008; Hyma and Fay, 2013). In northern Europe and Canada however, intensive sampling has yielded only *S. paradoxus* (Johnson *et al.*, 2004; Charron *et al.*, 2014; Kowallik *et al.*, 2015; Sylvester *et al.*, 2015; Leducq *et al.*, 2015). Without knowing the expected geographic distribution of the species, wild populations of *S. cerevisiae* remain challenging to find, hindering studies on its natural ecology and genetic diversity.

Experiments in the lab show that *S. cerevisiae* has a higher optimum growth temperature than *S. paradoxus* (Sweeney *et al.*, 2004; Salvadó *et al.*, 2011; Leducq *et al.*, 2014). Some aspect of seasonal temperature may therefore predict the differences in the geographic range of these species (Charron *et al.*, 2014; Leducq *et al.*, 2014). It seems unlikely that winter temperatures would be the best predictor of the differences in geographic distributions between the two species since they grow at similar rates at low temperatures (5-23°C; Sweeney *et al.*, 2004; Salvadó *et al.*, 2011). Furthermore, both *S. paradoxus* and *S. cerevisiae* strains isolated from North American oak trees show high tolerance to freezing and thawing (Will *et al.*, 2010). In contrast, *S. cerevisiae* strains grow much faster than *S. paradoxus* at temperatures over 30^°^C, and *S. cerevisiae* strains are typically able to grow at temperatures over 40°C whereas most *S. paradoxus* cannot (Liti *et al.*, 2009; Salvadó *et al.*, 2011). The optimum growth temperatures for both species (Sweeney *et al.*, 2004; Salvadó *et al.*, 2011) are also similar to maximum summer temperatures in Europe and North America (Hijmans *et al.*, 2005). Therefore, in this study we investigated summer temperature as a potential predictor of the geographic distributions of *S. cerevisiae* and *S. paradoxus*.

We surveyed for the presence of *S. cerevisiae*, *S. paradoxus*, and other budding yeast on oak trees in northern and southern Europe, where summer temperatures are especially low and high. As well as summer temperature, we considered other ecological variables that might be important in this habitat. For example, ancient oaks seem likely to harbour a much greater diversity of microbes than young trees, and thus we also collected trunk girth data as a proxy for tree age. We isolated wild *S. cerevisiae* only in southern Europe, and at a rate that was too low for a direct analysis of its distribution. Focusing instead on the distribution of its sister species, *S. paradoxus*, we detected associations between isolation rate, trunk girth and summer temperature, and used our model of these relationships to estimate the range of summer temperatures where *S. paradoxus* is predicted to be most abundant on oak trees. Using our estimated optimal temperature range for *S. paradoxus* and a laboratory estimate of the difference in temperature preference for woodland *S. cerevisiae* and *S. paradoxus* strains (Sweeney *et al.*, 2004), we predicted the worldwide geographic distributions of optimal summer temperatures for both species. In order to test our predictions, we compiled a dataset of sampling locations and genotype information that includes hundreds of *S. cerevisiae* as well as *S. paradoxus* isolates from previous studies (Liti *et al.*, 2009; Zhang *et al.*, 2010; Kuehne *et al.*, 2007; Leducq *et al.*, 2014; Naumov *et al.*, 1997; Cromie *et al.*, 2013; Wang *et al.*, 2012; Almeida *et al.*, 2015, and references therein). We show that the geographic distribution of *S. paradoxus* and wild *S. cerevisiae* is consistent with the potential ranges that we predict based on their optimal temperatures. We discuss the implications of our results for future field sampling and research into the ecology and evolutionary genetics of these and other yeast species.

## Materials and Methods

### Isolation of yeasts from fruit and oaks

Between September 2006 and November 2011, we collected 812 environmental samples from oak trees (UK, France and Greece), fruiting fig trees (Portugal and Greece), vineyard grapes (UK) and garden grapes (Greece) (Table 1, Table 2, Figure 1). The substrates tested for oak were mostly bark (*n* = 618), but a small number of soil samples (*n* = 15) were also collected at the base of some oak trees. The substrates tested for fig and grape were mostly fruit (*n* = 84 and *n* = 53, respectively), but also include fig bark (*n* = 9), grape bark (*n* = 21) and grape must (*n* = 12).

**Table 1:**
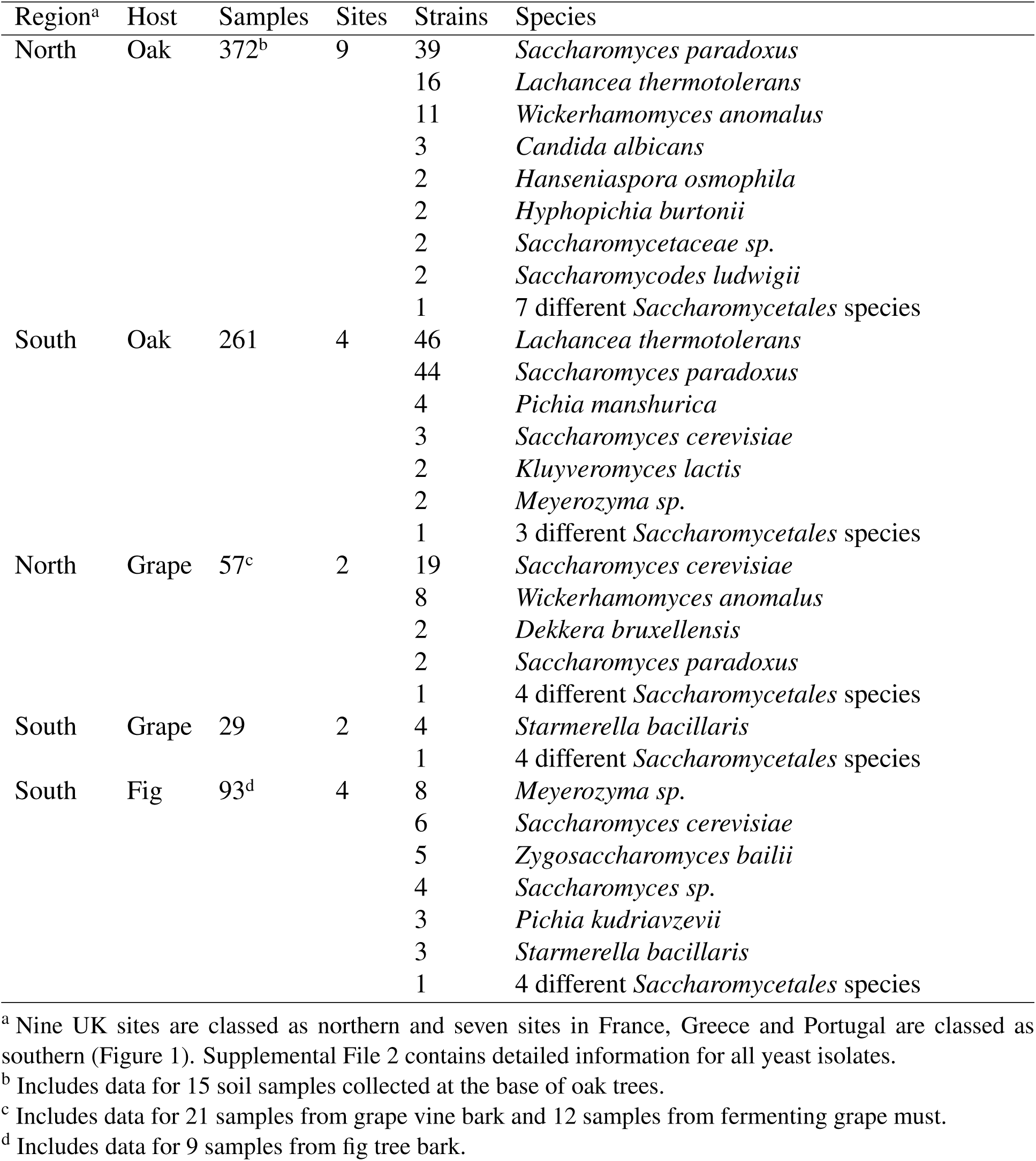
Yeast species isolated from oaks and fruits in northern and southern Europe

**Table 2:** Isolation frequencies of S. cerevisiae and S. paradoxus from oak bark

| Country | Site | Location | Trees<sup>a</sup> | Samples | Mean<br>$T_{max}$<sup>b</sup> | Mean<br>girth<sup>c</sup> | *Sc* | *Sp* | *Sp*<br>freq.<sup>d</sup> |
| --- | --- | --- | --- | --- | --- | --- | --- | --- | --- |
| U.K. | 1 | Brockholes Wood | 15 | 131 | 19.6 | 1.5 | 0 | 10 | 0.08 |
|  | 2 | Chorlton | 1 | 1 | 21.3 | 1.1 | 0 | 0 | 0.00 |
|  | 3 | Ladybower Wood | 4 | 32 | 19.6 | 2.3 | 0 | 7 | 0.22 |
|  | 4 | Tatton Park | 2 | 5 | 20.1 | 4.0 | 0 | 1 | 0.20 |
|  | 5 | Earlham Park | 2 | 3 | 20.9 | 6.8 | 0 | 1 | 0.33 |
|  | 6 | Fritham, New Forest | 15 | 60 | 21.3 | 3.3 | 0 | 7 | 0.12 |
|  | 7 | Ocknell, New Forest | 15 | 59 | 21.4 | 1.5 | 0 | 4 | 0.07 |
|  | 8 | Davenport Vineyard | 6 | 28 | 21.4 | 1.3 | 1 | 1 | 0.04 |
|  | 9 | Plumpton Vineyard | 6 | 24 | 21.6 | 1.3 | 0 | 3 | 0.12 |
| France | 10 | Montbarri, Bédarieux | 15 | 59 | 28.0 | 0.8 | 1 | 9 | 0.15 |
| Greece | 11 | Taxiarchis | 15 | 60 | 27.3 | 0.8 | 0 | 20 | 0.33 |
|  | 12 | Pyrgadikia | 15 | 82 | 30.9 | 1.4 | 2 | 14 | 0.17 |
|  | 13 | Parnitha | 15 | 60 | 29.7 | 1.1 | 0 | 1 | 0.02 |

<sup>a</sup> Includes data for 22 trees that were excluded from generalised linear models because of missing data for tree trunk girth (see Methods).
<sup>b</sup> Average of the daily maximum temperature in the hottest month of the year (°C). Weighted means are shown in cases where $T_{max}$ of trees differ within a site.
<sup>c</sup> Weighted mean trunk girth (m), weighted by the number of bark samples per tree.
<sup>d</sup> For each site, the number of *S. paradoxus* isolates / number of samples.

**Figure 1:**
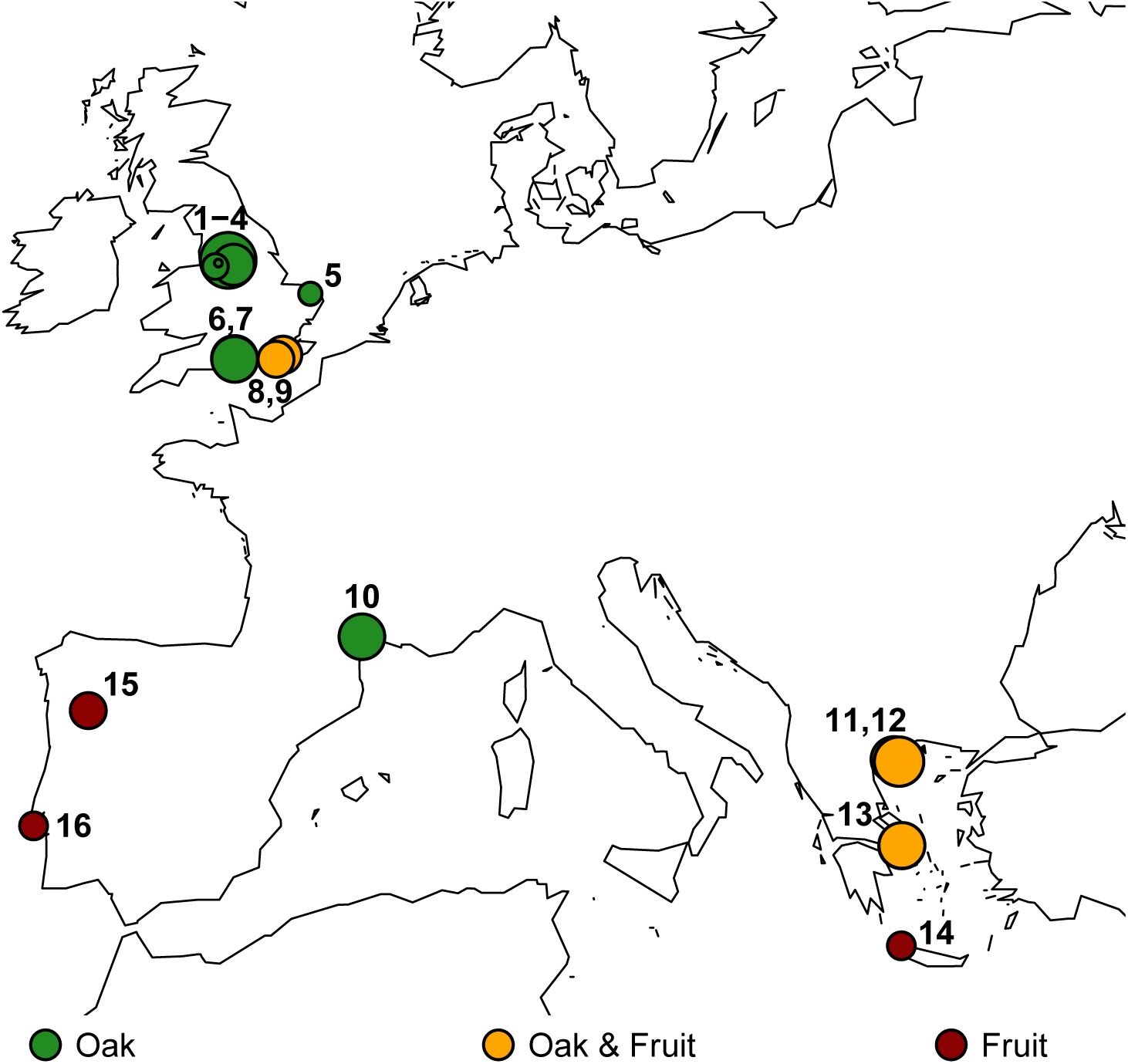
Sample collection sites for yeast strains isolated in this study. Circles are scaled by the natural log of the sample size. Numbers correspond to sites with oak trees in Table 2. No oak trees were sampled at field sites 14-16, and thus these sites were not included in Table 2.

Host plants were photographed and longitude and latitude were recorded in WGS84 format (https://github.com/bensassonlab/yeastecology/). Oak trees were classified as *Quercus robur, Q. petraea, Q. pubescens, Q. virgiliana, Q. frainetto* and *Q. ilex* using field guides (Sutton, 1990; Fitter and More, 2002). As an indicator of oak tree age, we measured trunk girth approximately 1m above the base of the tree. A number of the oak trees sampled were coppiced, and in these cases oak girth measurements taken from a single trunk underestimate the age of trees relative to uncoppiced trees. Using photographs of each tree, we treated trunk girth as missing data for 20 trees that were either coppiced or for which we could not determine coppicing status. No girth measurements were taken for an additional two trees sampled. In total, trunk girth data was missing for 22 trees out of 126 in our final statistical model.

Using sterile technique, environmental samples were collected from each host plant, stored in tubes for up to a week at room temperature, and weighed upon return to the laboratory. All samples were then incubated for at least two weeks in a liquid medium containing chloramphenicol and 7.6% ethanol that enriches for *Saccharomyces* (Sniegowski *et al.*, 2002). Most samples were incubated at 30^°^C, but 16 pilot samples were incubated at 10^°^C, and 18 at 25^°^C. Aliquots from 7.6% ethanol enrichment medium were streaked onto selective plates with a sole carbon source of methyl-*α*-D-glucopyranoside (Sniegowski *et al.*, 2002), and if weak yeast-like growth was seen on selective plates, then we also streaked from the 7.6% ethanol enrichment medium onto yeast extract peptone glucose (YPD) agar plates.

For each of the yeast-containing environmental samples, we picked multiple colonies from selective or YPD plates, pooled them in a single YPD liquid culture, and grew these pooled cultures to stationary phase. An aliquot of the pooled colony YPD liquid culture was preserved in 15% glycerol at -80°C, while the rest was used for DNA extraction. This pooled DNA was tested for the presence of our target species, *S. cerevisiae* and *S. paradoxus*, with species-specific PCR primers. In parallel, for every environmental sample that had yeast-like colonies on the original plates, we also picked a single colony into YPD liquid medium, preserved an aliquot of this single colony YPD culture, and identified the yeast species present. If tests on pooled DNA showed that an environmental sample contained *S. cerevisiae* or *S. paradoxus*, but the single colony culture contained a different species, then we plated the pooled culture and tested more individual colonies from this or from the original plate until we isolated *S. cerevisiae* or *S. paradoxus*. By testing both pooled samples and single colony cultures, it was possible to detect *S. cerevisiae* or *S. paradoxus* when other species were also present, as well as to detect *S. cerevisiae* and *S. paradoxus* in the same samples. As a result, we occasionally isolated *S. cerevisiae* or *S. paradoxus* with other yeast species from single environmental samples (8 out of 812 samples).

### Identification of yeast species

DNA was extracted from yeast using the Promega Wizard^®^ Genomic DNA purification kit, according to the manufacturer’s instructions for yeast, except that only 75 units of lyticase (Sigma) were typically used in an overnight incubation at 37°C. Conditions for PCR and DNA sequencing were as described in Bensasson (2011). DNA sequencing reads from PCR products were assembled using the Gap4 shotgun assembly tool of Pregap4 version 1.6-r (Bonfield *et al.*, 1995). Base accuracies were estimated by Pregap4 using its logarithmic (phred) scale. Consensus sequences were all exported from Gap4 (version 4.11.2-r.) in fasta format. Low quality consensus base calls were defined as those with a phred-scaled quality below q40, and were masked in the consensus sequence as “N”. Most DNA sequences (*n* = 300) had more than 200 high quality bases and fewer than 100 low quality bases and were submitted to NCBI [KT206983-KT207282]. A further 71 DNA sequences did not meet GenBank submission criteria, because they were technical replicates, were less than 200 bases long or contained more than 100 Ns, but were of sufficient quality for species identification and are available at https://github.com/bensassonlab/yeastecology/.

We used rapidly evolving centromeres (CEN6, CEN9 and CEN15) to identify *S. cerevisiae* and *S. paradoxus* strains (Bensasson *et al.*, 2008), and rDNA (18SrRNA-ITS1-5.8SrRNA-ITS2-25SrRNA) to identify other yeast species. All DNA samples were tested with primers specific to *Saccharomyces* CEN6, one *S. cerevisiae*-specific primer pair and one *S. para-doxus*-specific centromere primer pair (CEN6, CEN9 and CEN15; Bensasson, 2011; Supplemental file 1). In cases where PCR products were amplified using species-specific CEN primers, we sequenced at least one species-specific PCR product. All other DNA samples were tested using generic rDNA PCR primers (Supplemental File 1) and at least one rDNA sequence was generated for every isolate. We designed generic rDNA primers using primer3 (http://primer3.sourceforge.net/) that would anneal to all known Saccharomycetales rDNA sequences (in NCBI, June 2007), including 15 different Debaryomycetaceae and Saccharomycetaceae species.

Each isolate was then classified on the basis of the similarity of its centromere or rDNA to known yeast species using NCBI BLAST (https://blast.ncbi.nlm.nih.gov/). Every DNA sequence was queried against the nucleotide collection (nr/nt, date: August 28th, 2015) database restricted to the Ascomycota (taxid:4890), excluding a strain with *Lachancea thermotolerans* rDNA sequence that was classified as *S. paradoxus* in GenBank (Entrez Query “NOT LL12_027”). Searches were performed using the blastn algorithm (version 2.2.32+), with an expect threshold of 0.001, and no filtering for low complexity regions. Blast output was parsed using a custom perl script to extract the species names for hits with the highest blast score, and to assign species given a set of species name synonyms defined in the NCBI taxonomy (Supplemental File 2. For most yeast isolates (*n* = 247), species assignment was unambiguous; all hits with the highest BLAST score belong to only a single species (sometimes with multiple synonyms), and we assumed this was the species isolated. For a few strains (*n* = 17), DNA sequence had equal BLAST scores for multiple species, and in these cases we could only assign species to genus or higher taxonomic levels.

### Statistical analysis

All statistical and graphical analyses were conducted in R, version 3.1.1. Maps were drawn using the raster (version 2.3-40) and maps (version 2.3-9) packages using summer temperature (T*_max_*) data from the WorldClim dataset version 1.4 (1950-2000, release 3, http://www.worldclim.org) at 10 arc-minute (Figure 4) or 30 arc-second (approximately 1km) resolution (Figure 5, Supplemental files 1 and 4) (Hijmans *et al.*, 2005). T*_max_* was estimated using raster for every host plant from a single pixel at 30 arc-second resolution. T*_max_* in the WorldClim dataset is the daily maximum temperature, averaged over the hottest month of the year (Robert Hijmans, personal communication).

Using a generalised linear model (GLM) with binomial errors, we modelled *S. paradoxus* isolation frequency by setting the proportion of bark samples with *S. paradoxus* from an oak tree as the response variable. The initial model included four explanatory variables and all their possible interactions: (i) trunk girth (in metres) as a continuous variable; (ii) T*_max_* (in °C×10) as a continuous variable estimated from a single pixel at 30 arc-second resolution given the longitude and latitude of each tree; (iii) a three level factor describing oak type as robur-like (the northern *Q. robur* or *Q. petraea*), frainetto-like (the southern *Q. frainetto, Q. pubescens* or the intermediate *Q. virgiliana*) or the outgroup species *Quercus ilex*; (iv) a continuous variable describing the frequency of non-*S*. *paradoxus* yeast species isolation (the number of other yeast species isolated divided by the number of samples collected for each tree). This initial model was simplified by subtracting terms in a stepwise manner starting from the highest order terms and testing whether each subtraction resulted in a worse model using χ^2^ tests as recommended in Crawley (2005). The three-level factor for oak type was then further simplified to two levels and nested models were again compared using *χ*^2^ tests following the principles for model simplification by contrasts described in Crawley (2005).

Both the initial and final models showed expected levels of deviance given the number of degrees of freedom (final model, residual deviance=75, *d.f.*= 98). Cook’s distance analysis was also used to identify the trees with the highest influence on the parameter estimates of the model. As a control we investigated the effects of each of these data points on the analysis, and found the removal of single data points did not qualitatively change the final model. To control for the possibility that a single site in southern Europe affects our conclusions, we investigated the effects on the analysis of dropping all data for one southern field site at a time. In all cases, we observed all the same statistically significant effects (*P* < 0.04), and visualisation of the effects showed no qualitative difference from the results shown in Figures 2 and 3.

**Figure 2:**
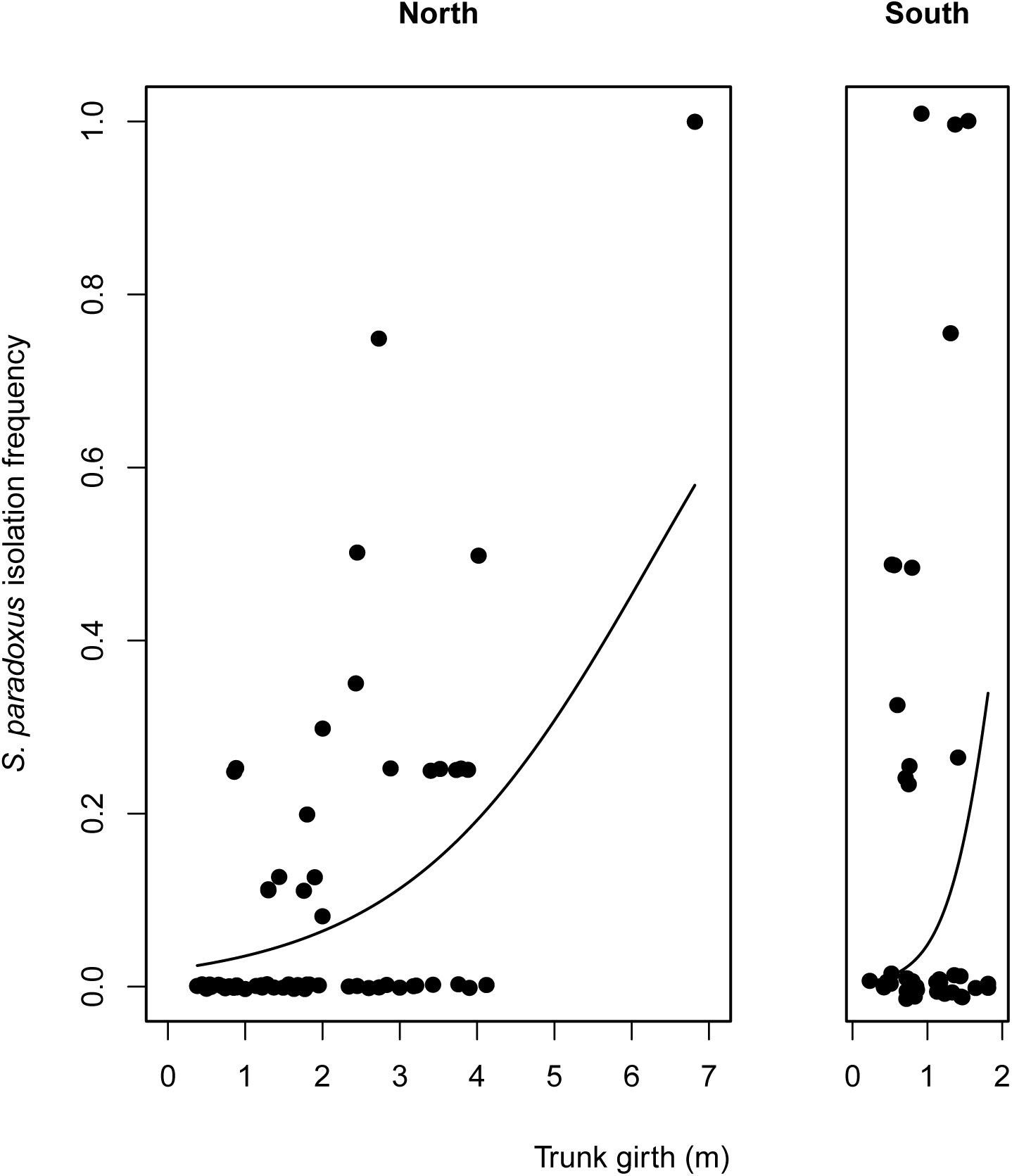
*S. paradoxus* isolation frequency increases with trunk girth. Points show the observed isolation frequencies for 104 trees from northern (UK) and southern Europe (France and Greece). For each tree, we estimated the frequency of *S. paradoxus* isolation as the number of pieces of bark yielding *S. paradoxus* divided by the number of pieces of bark sampled. Points are clustered around discrete frequencies because in most cases the number of pieces of bark sampled was four. We therefore used jitter to allow better visualisation of data. Lines show the probability of isolating *S. paradoxus* estimated from the final GLM assuming median summer temperatures in northern (T*_max_* =21.3°C) and southern Europe (T*_max_*=28.6°C)

**Figure 3:**
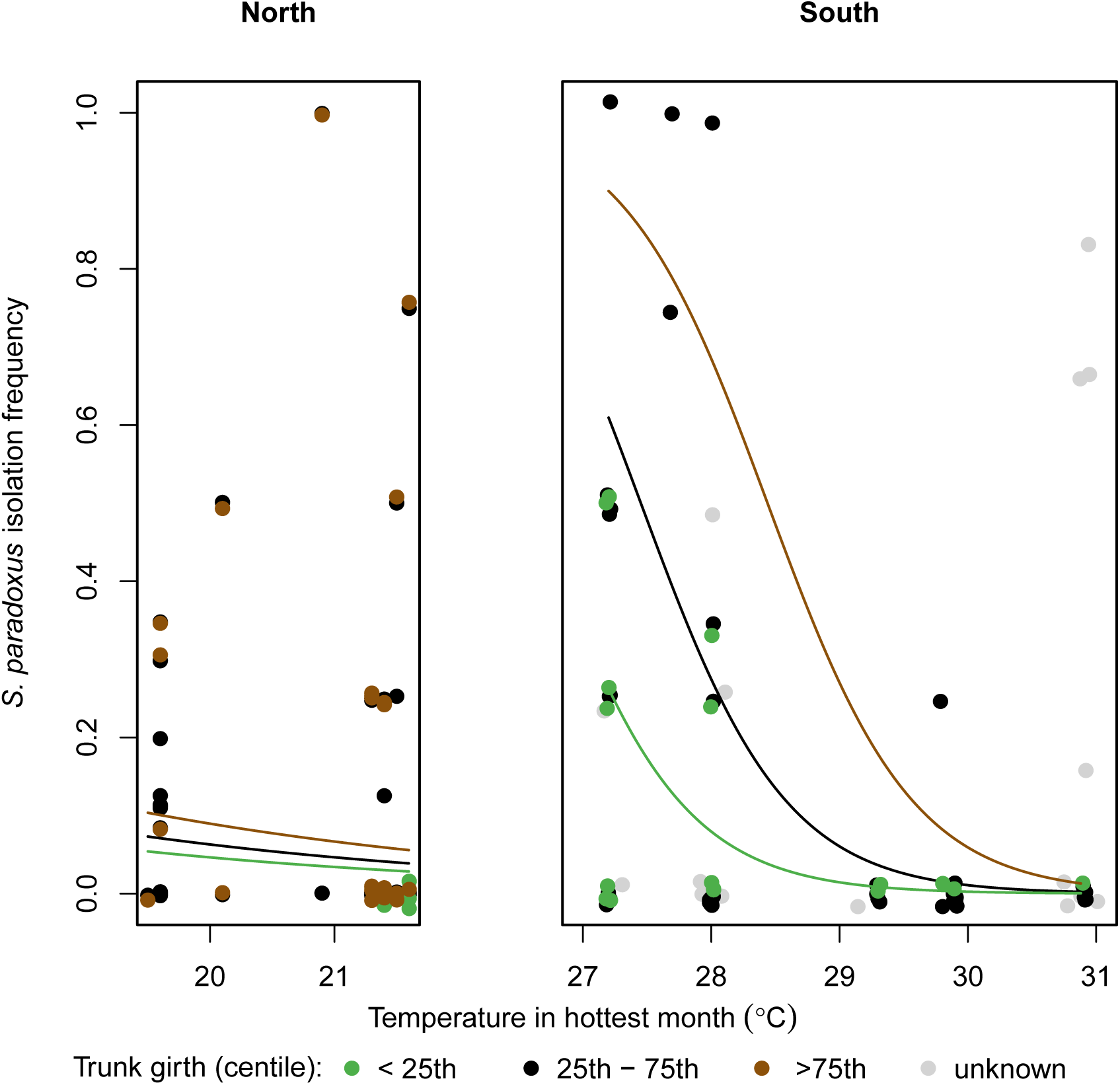
The effects of temperature in the hottest month *S. paradoxus* isolation frequency. *S. paradoxus* isolation frequency is estimated as the proportion of bark samples from each tree with *S. paradoxus;* more specifically, the number of *S. paradoxus* isolates for a tree divided by the total number of bark samples obtained for that tree. Points show the distribution of the data, including points for which no trunk girth data are available (grey, see Methods). Jitter was used to better display overlapping points. Lines show the predicted probability of isolating *S. paradoxus* and are estimated from the final generalised linear model given lower (0.8m), median (1.3m), and upper (1.9m) quartile measurements of tree trunk girth (green, black and brown respectively).

### Worldwide presence and absence data for S. *paradoxus* and S. *cerevisiae*

In order to test whether *S. cerevisiae* and *S. paradoxus* have been isolated from locations with summer temperatures within the optimum ranges that we predict, we needed sample location and genotype information for a large number of strains. Sampling locations have been mapped for thousands of yeast strains from many species that have been deposited in the Centraalbureau voor Schimmelcultures collection (Robert *et al.*, 2006; Kurtzman *et al.*, 2015). This resource is not available for download however, and does not provide genotype information, which we need in order to distinguish wild from human-associated *S. cerevisiae* strains. Location information has been mapped together with genotype information for *S. paradoxus* (Boynton and Greig, 2014), but not for *S. cerevisiae.*

Therefore we collated site location information together with genotype information from previous studies on *S. cerevisiae* (Zhang *et al.*, 2010; Wang *et al.*, 2012; Cromie *et al.*, 2013; Almeida *et al.*, 2015) and *S. paradoxus* (Naumov *et al.*, 1997; Kuehne *et al.*, 2007; Liti *et al.*, 2009; Zhang *et al.*, 2010; Leducq *et al.*, 2014). No data for *S. paradoxus* strains isolated in this study that were used in the construction of our statistical model were included in this validation dataset. Site location and genotype information for *S. cerevisiae* strains isolated as part of this study were included, because no information for these strains was used to generate the model. The criteria for including data from a study were that it provided genotype information for many strains (that are not already included in a larger study) and it included strains isolated from substrates that are not wine or vineyard grapes. In most previous studies, latitude and longitude information was not included in site descriptions. We therefore used site descriptions as search terms in Google Maps. Where site descriptions map to a large region, we used latitude and longitude coordinates from the estimated centre of that region. Data for yeast strains with site descriptions that did not allow location within 100-200 km were excluded (for example, strains from unknown locations or with their origin described as “Europe”). We also excluded strains isolated from wine or vineyard grapes, because we expect that their distribution is affected by human activity (Fay and Benavides, 2005). *S. cerevisiae* was also recorded as absent from several sites where surveys of over 100 bark samples yielded no *S. cerevisiae:* site 1 from this study (Table 2), Charron *et al.* (2014), Johnson *et al.* (2004) and Kowallik *et al.* (2015).

T*_max_* was estimated for every isolate using the raster package from a single pixel at 30 arc-second resolution. For collection sites that occur at locations with summer temperatures outside the range that we predict with our statistical model, we estimated the distance to regions that are within the expected range. The region in which such sites occurred were visualised using the raster and maps packages in R, and the distance (in kilometres) was estimated using the sp package in R (version 1.1-1).

## Results

### Variation in the geographic distribution of yeast species

We conducted a field survey with the aim of isolating yeast species from the *Saccharomyces sensu stricto* genus, and isolated 264 yeast strains from 812 European oak, fig, and grape samples (Table 1, Figure 1, Supplemental File 3). These strains are from at least 26 different yeast species across the order Saccharomycetales, including 5 different yeast families: Saccharomycetaceae, Saccharomycodaceae, Debaryomycetaceae, Phaffomycetaceae, and Pichiaceae (Supplemental File 2). Although it is rarely isolated in natural environments (Tanghe *et al.*, 2005; Lachance *et al.*, 2011; Maganti *et al.*, 2011), we isolated three strains of the human commensal and pathogen, *Candida albicans* from ancient oak trees in northern Europe (site 6 in Figure 1 and Table 2, Supplemental File 1). *C. albicans* has only rarely been isolated away from mammals (Tanghe *et al.*, 2005; Lachance *et al.*, 2011; Maganti *et al.*, 2011), and the existence of wild populations of *C. albicans* on north European trees could potentially explain the hitherto puzzling maintenance of aquaporin genes that confer freeze tolerance in *C. albicans* (Tanghe *et al.*, 2005).

The most commonly isolated *Saccharomyces* species was *S. paradoxus*, which we isolated mostly from oak bark and from soil at the base of oak trees (83 out of 633 samples, Table 1). We isolated *S. cerevisiae* strains from 25 out of 179 fruit, fruit tree bark and grape must samples, but relatively few from oak-associated samples (4 out of 633, Table 1). In addition, we isolated a single strain of *S. kudriavzevii* from oak bark in Greece (site 12, Figure 1) as well as four strains of a *Saccharomyces sensu stricto* species from figs at the same site that we could not identify to the species level using our methods (Table 1). The greater prevalence of *S. cerevisiae* on fruit trees relative to oaks could however be an effect of geography and human influence, because fruit trees were only sampled in the far south of Europe or in vineyards (Figure 1, Table 2). Indeed, when we controlled for the effects of geography by considering only sites where *S. cerevisiae* was present, we saw very similar isolation rates from fruit, fruit tree bark and oak bark (Supplemental File 1). Others have also observed similar or lower isolation rates from fruit relative to woodland substrates (Wang *et al.*, 2012), and this finding lends support to the proposal that *S. cerevisiae* is not adapted to fruit (Goddard and Greig, 2015).

In the UK, we isolated 39 *S. paradoxus* from 372 oak bark and soil samples (Table 1). This isolation rate (10%) is similar to that previously reported for *S. paradoxus* both in the UK (Johnson *et al.*, 2004) (28 isolates from 344 oak bark samples, Fisher’s exact test, *P* = 0.3) and Pennsylvania, USA (Sniegowski *et al.*, 2002) (8 out of 79 oak bark and soil samples, Fisher’s exact test, *P* = 1). In contrast, we isolated fewer *S. cerevisiae* from oak samples in the UK (1/372) than Sniegowski *et al.* (2002) did from oak trees in Pennsylvania (10/79; Fisher’s exact test, *P* = 2 × 10^−7^), even though we used the same enrichment culturing method. The fact that we were able to reproduce the *S. paradoxus* isolation rate, but not the *S. cerevisiae* isolation rate (Sniegowski *et al.*, 2002), suggests a geographic difference in the distribution of *S. cerevisiae* relative to *S. paradoxus*, with a lower abundance of *S. cerevisiae* in the UK than in Pennsylvania.

Analysis of all 264 strains isolated from all 812 European samples suggests that there are also differences in the geographic distributions of other yeast species within Europe (Table 1). In general, we were able to isolate and identify more yeast strains from southern than from northern European oak bark (104/261 compared to 84/372, Fisher’s exact test, *P* = 4 × 10^−6^). This effect is especially strong for *Lachancea thermotolerans,* a yeast common in oak bark (Sampaio and Gonçalves, 2008; Sylvester *et al.*, 2015), which is more common in southern (46 out of 261) than in northern oak bark and soil samples (16/372; Fisher’s exact test, *P* = 4 × 10^−8^, Table 1). Previous studies have shown enrichment culturing at different temperatures (10°C compared to 22-30°C) results in the isolation of different yeast species (Sampaio and Gonçalves, 2008; Sylvester *et al.*, 2015). Therefore the bias toward southern yeast distributions might simply be a consequence of the temperature we use for enrichment culturing (25-30°C). However, it is not a universal rule that all yeast species have higher isolation rates in southern versus northern locations. Notably, *Wickerhamomyces anomalus,* a food spoilage yeast that can also contribute to wine aroma (Passoth *et al.*, 2006), was common in northern oak (11 out of 372 bark and soil samples) and fruit, but was absent from southern oak bark samples (0/261; Fisher’s exact test, *P* = 0.004) and fruit (Table 1).

### Trunk girth and summer temperature can explain differences among oaks in *S. paradoxus* abundance

The original aim of this study was to model the ecological factors affecting the prevalence of *S. cerevisiae* in woodlands, but consistent with other studies on northern European sites (Johnson *et al.*, 2004; Kowallik *et al.*, 2015), we were unable to isolate many *S. cerevisiae* strains from European oaks. Instead, we focused our modelling efforts on its closest relative *S. paradoxus*, which was the most commonly isolated species in this study (Tables 1 and 2). For these analyses we used data for 78 strains of *S. paradoxus* isolated from 126 oak trees resulting from a total of 604 oak bark samples (Table 2). An average of 4.8 pieces of bark were collected from each tree, and in most cases (87 trees), we collected exactly 4 pieces per tree. To reduce potential variation resulting from experimental procedures, we excluded pilot data for 14 oak bark samples that were incubated at 10°C during enrichment culturing and 15 soil samples collected at the base of oak trees. Preliminary analysis showed that isolation rates are not affected by collection month and bark sample weight in this study (Supplemental File 1), and therefore these variables were not included in our final model.

Lab studies suggest that *S. cerevisiae* and *S. paradoxus* have different temperature preferences for their optimal growth (Sweeney *et al.*, 2004; Salvadó *et al.*, 2011) and also differ in their tolerance of high temperatures (Liti *et al.*, 2009). Therefore, we asked whether summer temperature (T*_max_*) can predict the distribution of *S. paradoxus,* in conjunction with other variables that could affect the prevalence of yeast on oak trees, such as host species or tree age. Because other yeast species could potentially outcompete *S. paradoxus* in culture and affect our estimation of its isolation rate, we also consider the presence of other yeast species isolated from each tree in our analysis. Using trunk girth as a proxy for tree age, and binning tree species into three groups (robur-like, frainetto-like, and *Q. ilex*; see Methods), we constructed a generalised linear model (GLM) to test whether the frequency of *S. paradoxus* isolation from an oak tree can be predicted by four explanatory variables (i) trunk girth, (ii) summer temperature, (iii) host tree type, and (iv) isolation frequency of other yeast species.

After standard model simplification (Crawley, 2005), we found that the presence of other yeast species does not affect the number of *S. paradoxus* isolated (GLM, -0.02% deviance, *d.f.* = 1, *P* = 0.9). This suggests that competition among yeast during our isolation procedure does not substantially affect the rate or pattern of *S. paradoxus* isolation. However, all three other explanatory variables are important for predicting numbers of *S. paradoxus* isolated from oak trees. We also found that a simpler final model where oaks are classed as northern or southern is not worse than the model describing three host types (GLM, -2% deviance, *d.f.* = 3, *P* = 0.4). This suggests that more general differences between northern and southern European field sites can explain differences in *S. paradoxus* yield better than host tree type.

The final GLM explains 42% of the deviance among trees in *S. paradoxus* isolation frequency in terms of tree trunk girth, summer temperature, and whether a site is northern or southern. Trunk girth is an important predictor of *S. paradoxus* isolation frequency, which if dropped leads to a much worse model fit (GLM, -21% deviance, *d.f*. = 2, *P* = 1 × 10^−6^). Indeed, if we remove trunk girth data from the analysis, we find that none of the other significant effects in the model would have been detected, suggesting that host tree age is a crucial factor to consider in order to discover variables that are relevant to yeast ecology. As trunk girth increases, *S. paradoxus* isolation frequency increases in northern and southern Europe (Figure 2). The positive association between trunk girth and the presence of *S. paradoxus* suggests that old oak trees harbour more *S. paradoxus*.

The best predictor of the *S. paradoxus* isolation frequency for a tree was whether it was from northern or southern Europe. Trees from southern Europe yielded more *S. paradoxus* isolates, even though we sampled more trees and larger trees from northern Europe (Table 2, Figure 3). This effect is especially clear in Figure 3 from the low isolation frequency of *S. paradoxus* that the model predicts in northern Europe compared to the high frequency expected at temperatures around 27-28^°^C in southern Europe. There is also a difference between northern and southern trees in the effect of trunk girth on *S. paradoxus* isolation frequency (GLM, -6% deviance *d.f*. = 1, *P* = 0.004). More specifically, the numbers of *S. paradoxus* isolated from southern oaks increased more steeply with increasing trunk girth than they did from northern oaks (Figure 2).

In southern Europe, we also observe a negative relationship between *S. paradoxus* abundance and summer temperature, whereas there is no such effect in the north (GLM, -9% deviance, *d.f*. = 1, *P* = 0.0006, Figure 3). This suggests that the hottest field sites in southern Europe (T*_max_*, 28-31°C) are hotter than the optimum habitat for *S. paradoxus,* which is consistent with laboratory observations of suboptimal growth for most strains of *S. paradoxus* at temperatures over 30°C (Sweeney *et al.*, 2004; Salvadó *et al.*, 2011; Leducq *et al.*, 2014).

Figure 3 shows the predictions of the final model with all the variables of major effect combined. The low predicted *S. paradoxus* isolation frequency between 18 and 22°C suggests an optimum summer temperature for *S. paradoxus* that is higher than 22^°^C, whereas the negative association between T*_max_* and isolation rate between 28 and 31°C, suggests that the optimum is lower than 28^°^C. Thus, the optimum summer temperature for *S. paradoxus* appears to be between 22 and 28^°^C.

### Summer temperature can predict the worldwide distribution of wild *S. paradoxus* and *S. cerevisiae* populations

Our analysis of oak bark samples collected from thirteen European sites in the UK, France and Greece (Table 2, Figure 3) suggests that the optimum summer temperature (T*_max_*) for *S. paradoxus* lies between 22 and 28^°^C, but that this species is also found at lower abundances between 18 and 31^°^C (Figure 3). We tested the predictions of our model by mapping the global distribution of this thermal optimum, and comparing it to sites where *S. paradoxus* has been reported in previous studies (Naumov *et al.*, 1997; Kuehne *et al.*, 2007; Liti *et al.*, 2009; Zhang *et al.*, 2010; Leducq *et al.*, 2014). Virtually all the *S. paradoxus* strains that we mapped from other studies (244 out of 246) fall within our predicted range of optimum summer temperatures between 18 and 31°C (Figure 4A). Indeed, 75% of these *S. paradoxus* strains map to locations where T*_max_* is between 22 and 28°C, and 95% occur between 20 and 30°C. We identified only two strains that could fall outside the T*_max_* range of 18 to 31°C. One was from Tashkent in Uzbekistan (Naumov *et al.*, 1997), a site that we approximately mapped to the centre of Tashkent (with a T*_max_* of 36°C). This approximate mapping is within 30 km of high elevation regions that have a lower summer temperature (T*_max_* of 28°C), which is within our predicted optimum range. The other exception was a strain of *S. paradoxus* isolated from insect excrement (from Salem, MO, USA, 32°C T*_max_*; Leducq *et al.*, 2014), collected over 200km from locations with temperatures within the predicted range. This was one of only few animal-associated *S. paradoxus* strains (8 out of 246 strains), and the unusual location of this sample may possibly have arisen by insect mediated transport from a location with expected summer temperatures.

**Figure 4:**
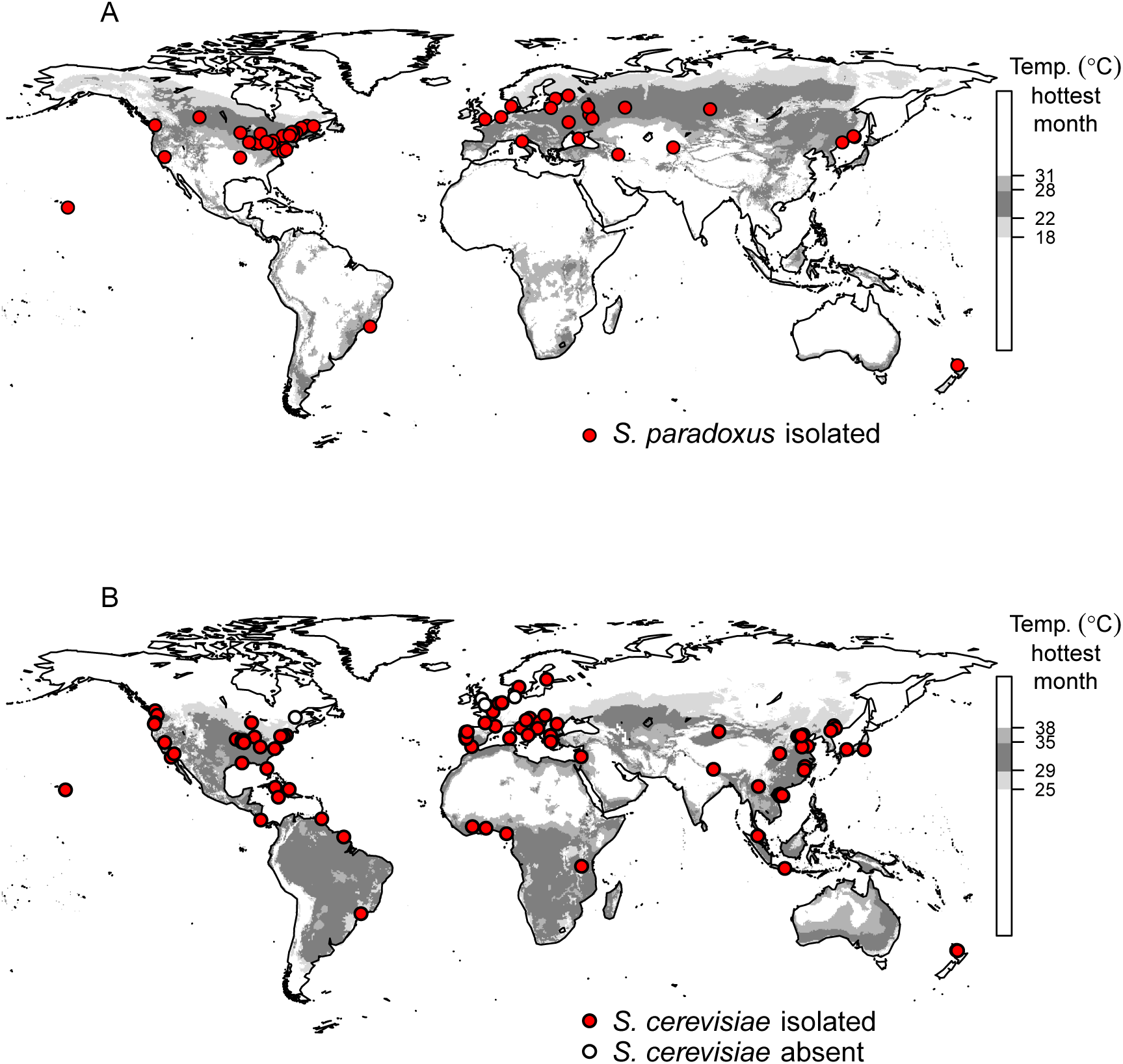
Global distribution of the predicted optimum temperature range for (A) *S*. *paradoxus* and (B) *S*. *cerevisiae.* Optimum temperatures for *S. paradoxus* are estimated from Figure 3, and for *S. cerevisiae* we assume the optimum is approximately 7°C higher than that of *S. paradoxus* (Sweeney *et al.*, 2004). Red circles show the approximate origin of strains published in large genotyping studies (Liti *et al.* (2009); Zhang *et al.* (2010); Kuehne *et al.* (2007); Leducq *et al.* (2014); Naumov *et al.* (1997); Cromie *et al.* (2013); Wang *et al.* (2012); Almeida *et al.* (2015) and references therein). Location and genotype (Almeida *et al.*, 2015) information from this study is included for *S. cerevisiae* strains but not for *S. paradoxus*, because data for *S. paradoxus* were used to generate our predictions. White circles show locations where surveys of over 100 bark samples yielded no *S. cerevisiae* and are summarised from this study, Charron *et al.* (2014), Johnson *et al.* (2004) and Kowallik *et al.* (2015).

Ideally, we would like to map the worldwide distribution of the model eukaryote, *S. cerevisiae*. We can make progress towards this goal by combining our results from *S. paradoxus* with the finding by Sweeney *et al.* (2004) that in the laboratory, *S. cerevisiae* from oak trees grow optimally at roughly 7^°^C higher temperatures than *S. paradoxus*. We use the estimate of the species difference in temperature preferences by Sweeney *et al.* (2004), because this study uses a large number of *S. cerevisiae* and *S. paradoxus* strains from the same oak habitat, with growth profiles that are typical for their species (see Supplemental File 1 for a full discussion). In order to predict the potential geographic range of *S. cerevisiae,* we therefore added 7^°^C to our climate envelope model for *S. paradoxus* to generate a global distribution map based on predicted optimum temperatures for *S. cerevisiae* (Figure 4B). The potential range that we predict for *S. cerevisiae* is mostly subtropical or tropical and different from the prediction of a temperate distribution for *S. paradoxus* (Figure 4). Indeed, the predicted worldwide range of *S. cerevisiae* is clearly more consistent with the distribution of *S. cerevisiae* isolates than that of *S. paradoxus* (Figure 4).

Human culture and transport of *S. cerevisiae* across the world has affected the distribution of this species (Fay and Benavides, 2005; Liti *et al.*, 2009; Wang *et al.*, 2012; Cromie *et al.*, 2013). Therefore, when testing the predicted distribution of optimum summer temperature for *S. cerevisiae,* we need to distinguish strains that are associated with human activity from wild strains. Strains associated with human activity, such as those cultured in breweries or vineyards, can potentially escape and survive in regions with otherwise unsuitable climates as feral strains, but these are likely to represent transient (sink) populations. The locations of sink populations do not accurately test the predictions of climate envelope models (Araújo and Peterson, 2012). Feral *S. cerevisiae* strains are expected to have genotypes associated with human activity, such as the genotype associated with wine production, or to be “mosaic” strains showing recent genomic admixture between natural populations (Fay and Benavides, 2005; Liti *et al.*, 2009; Wang *et al.*, 2012; Cromie *et al.*, 2013; Almeida *et al.*, 2015).

The majority of *S. cerevisiae* isolates (222 out of 301 strains) from most of the collection sites (71 out of 92 sites) that we were able to map worldwide, mapped approximately to locations with summer temperatures within the optimum range that we predict for *S. cerevisiae* (25-38^°^C). Almost half the collection sites outside our predicted range occur in Europe (10 out of 21 sites) where yeast sampling intensity is relatively high (Robert *et al.*, 2006; Kurtzman *et al.*, 2015). Figure 5 shows all the *S. cerevisiae* strains (*n* = 46) isolated from Europe with points coloured according to genotype. Two distinct genetic lineages of *S. cerevisiae* predominate within Europe (Cromie *et al.*, 2013; Almeida *et al.*, 2015); one is associated with humans and wine and another is associated with oak trees (Almeida *et al.*, 2015) and perhaps also olive trees (Cromie *et al.*, 2013). The vast majority of European *S. cerevisiae* with the wild genotype expected on oak trees (23 out of 26 strains) map to locations with summer temperatures within the range that we predict for *S. cerevisiae* (between 25 and 38°C, Figure 5). The three wild strains in Europe that we mapped to locations outside the predicted range of summer temperatures mapped to Mount Subasio in Italy and Jasenovo polje in Montenegro (Figure 5). The locations for both of these sites were mapped approximately, and both occur in mountain regions with expected summer temperatures at lower elevation (within 3km). In contrast, several European strains with human-associated genotypes (7 out of 20 strains) occur at sites that are far from the predicted summer temperatures for *S. cerevisiae* (200-1300km away). Many of these strains with human-associated genotypes were isolated from locations that suggest a recent association with humans or that they could represent transient populations: a vineyard tree, buttermilk, a fish’s gut, and soil at an agricultural college. It therefore appears that in Europe, *S. cerevisiae* strains that fell outside our predicted range were either rare strains with wild genotypes that were probably incorrectly mapped to higher elevations in mountain ranges, or more commonly human-associated *S. cerevisiae* that can occur at locations far from our predicted range (Figure 5).

**Figure 5:**
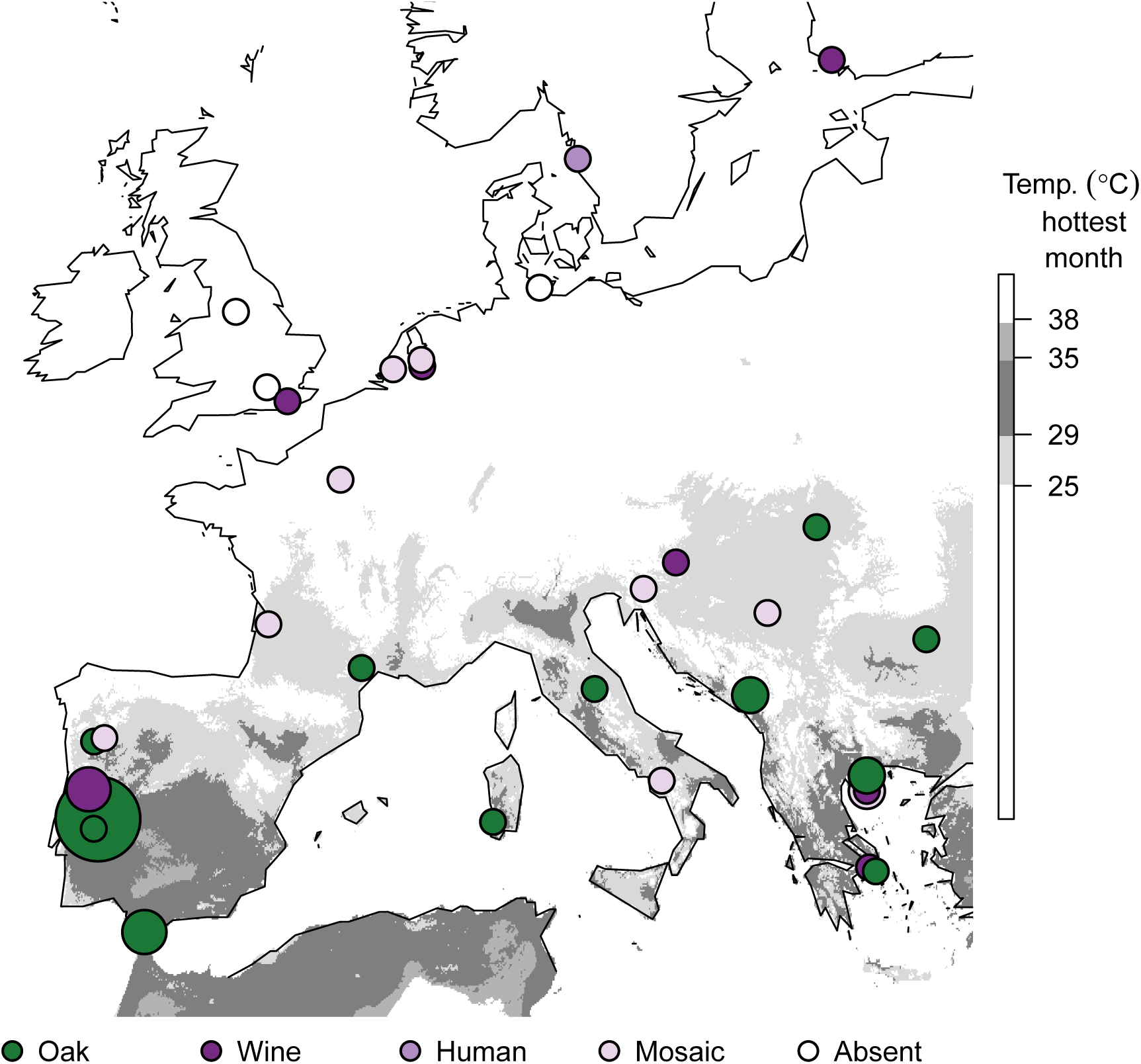
Only feral *S. cerevisiae* or those with mosaic genotypes occur outside the predicted optimal temperature range. The regions with average temperature in the hottest month where we expect *S. cerevisiae* are shaded in grey, assuming it correlates with a 7°C higher average temperature in the hottest month than *S. paradoxus* (Sweeney *et al.*, 2004). White points show the locations where over a hundred pieces of bark yielded no *S. cerevisiae* (this study, Johnson *et al.*, 2004; Kowallik *et al.*, 2015). The remaining points show the geographic sources of 46 *S. cerevisiae* strains isolated from various sources that include trees, soil, fruits and beer (but not including wine or grapes), and are coloured by genotype (see Results; data from Cromie *et al.*, 2013; Almeida *et al.*, 2015). Points are scaled by the square root of sample size and two points in Greece were repositioned slightly to so that all overlapping points are visible.

The patterns that we see in Europe are similar to those we see worldwide. *S. cerevisiae* strains have been isolated from soil, vine bark and buttercups in a New Zealand vineyard (Goddard *et al.*, 2010) outside the predicted range of summer temperatures (24°C, Figure 4B). These strains have genotypes similar to those of European rather than Asian *S. cerevisiae* (Cromie *et al.*, 2013) and thus may also represent vineyard-associated sink populations. Out of 122 *S. cerevisiae* strains with human-associated genotypes mapped worldwide, 38 strains occur at locations with summer temperatures that are lower than those we predict for *S. cerevisiae,* and 36 of these are more than 20km from locations with expected temperatures (Figure 5, Supplemental File 4). In contrast, the 41 out of 179 *S. cerevisiae* strains with wild genotypes outside the predicted range were much closer to locations within the predicted range than those with human-associated genotypes (Wilcoxon test, *P* = 9 × 10^−14^). All 41 wild *S. cerevisiae* strains that were out of range were mapped only approximately, and 40 of these mapped to mountain locations in Europe and China that were within 8km of the predicted range (median distance = 1km; Figure 5 and Supplemental File 1). The only exception of a strain with a wild genotype occurring far out of range was isolated from a flower in Seattle (T*_max_* 23°C, 84km from the nearest site within range; Cromie *et al.*, 2013). We therefore conclude that the distribution of wild *S. cerevisiae* strains is consistent with our predicted range.

In addition, our model correctly predicts most of the differences and similarities in the ranges of *S. cerevisiae* and *S. paradoxus*. The difference in the optimum summer temperatures illustrated in Figure 4 can explain the presence of *S. paradoxus* and the absence of *S. cerevisiae* in the UK (T*_max_* 20°C, this study; 23°C Johnson *et al.*, 2004), Canada (T*_max_* 25°C, Charron *et al.*, 2014) and northern Germany (T*_max_* 21°C, Kowallik *et al.*, 2015). Conversely, the optimum summer temperatures for the two species overlap between 25 and 31°C, where we might therefore expect their sympatry: for example, in the northern USA, parts of southern Europe, northern China, southeastern Brazil, South Africa, and southern Australia. In the northern USA (T*_max_* 30°C, Sniegowski *et al.*, 2002), and southern Europe at least (T*_max_* 31°C, Sampaio and Gonçalves, 2008; Table 2), these prediction are met.

## Discussion

By intensively sampling *S. paradoxus* from oak trees in northern and southern Europe (Figure 1, Supplemental File 3), we discovered associations between *S. paradoxus* isolation frequency, trunk girth (Figure 2) and summer temperature (Figure 3). Using the association of *S. paradoxus* with summer temperature in Europe, we predict regions where *S. paradoxus* and *S. cerevisiae* might occur worldwide (Figure 4). The worldwide distribution predicted by the optimum T*_max_* for *S. paradoxus* is consistent with the observed distribution of *S. paradoxus* isolations from previous studies (Boynton and Greig, 2014; Figure 4A, Supplemental File 4), and with the detection of a northern limit to its distribution in Canada (Charron *et al.*, 2014; Leducq *et al.*, 2015). Similarly, our predicted optimum summer temperature for *S. cerevisiae* could potentially explain the success or failure to isolate *S. cerevisiae* in previous studies (Figure 4B and Supplemental File 4; Johnson *et al.*, 2004; Charron *et al.*, 2014; Kowallik *et al.*, 2015), and why *S. cerevisiae* strains isolated outside this range often have human-associated or mosaic genotypes indicative of transient populations (Figure 5 and Supplemental File 4).

Population genetic analyses show that the genetic diversity of *S. cerevisiae* is exceptionally high in the tropics and subtropics of China (Wang *et al.*, 2012; Almeida *et al.*, 2015), and is unusually low in Europe (Almeida *et al.*, 2015). The genetic diversity of a population is expected to increase as its habitat area increases (Rauch and Bar-Yam, 2005). High genetic diversity of *S. cerevisiae* in China is therefore compatible with the larger potential habitat area we predict in east Asia (Figure 4B), while low genetic diversity within Europe is consistent with the restricted range predicted for *S. cerevisiae* in Europe (Figure 5). An alternative explanation for the high genetic diversity of *S. cerevisiae* in China is an east Asian origin for the species (Wang *et al.*, 2012; Almeida *et al.*, 2015). It is currrently unknown if other subtropical or tropical forest populations of *S. cerevisiae* have high genetic diversity since yeasts have been less intensively sampled from such regions (Robert *et al.*, 2006; Kurtzman *et al.*, 2015). Without further sampling in tropical and subtropical regions it is not possible to differentiate whether the higher diversity of *S. cerevisiae* in Asia reflects a greater habitat area or an Asian origin for *S*. *cerevisiae.*

Although our predictions fit well with the data currently available, this analysis represents only a starting point for understanding the ecological factors controlling the distribution of *S. paradoxus* and *S. cerevisiae.* In this study, we focused only on T*_max_* as a climate variable because laboratory experiments suggest a difference between *S. paradoxus* and *S. cerevisiae* in their growth at high temperatures (Sweeney *et al.*, 2004; Liti *et al.*, 2009; Salvadó *et al.*, 2011; Leducq *et al.*, 2014), but not at low temperatures (Sweeney *et al.*, 2004; Will *et al.*, 2010; Salvadó *et al.*, 2011). Different climate variables are highly correlated within Europe, and using only the field sites in this study (Table 2), we cannot distinguish the association of *S. paradoxus* isolation frequency with summer temperature from associations with other factors such as rainfall or winter temperature. Furthermore, our observation of a negative association between T*_max_* and *S. paradoxus* isolation frequency is based on analysis of data from only four independent field sites in southern Europe. Our conclusions would be strengthened by independent verification of the upper limit of the optimum T*_max_* for *S. paradoxus* from additional sites. Thus, while we conclude that summer temperature can predict the range of *S. paradoxus* and *S. cerevisiae*, we do not claim that summer temperature is the causal factor limiting the distribution of *Saccharomyces* species.

In the case of *S. cerevisiae*, our predictions are based indirectly on ecological findings for *S. paradoxus* and laboratory growth experiments from North American strains (Sweeney *et al.*, 2004). In using this laboratory estimate, we assume that the physiological response to temperature is fixed within species. However, the *S. paradoxus* strains used by Sweeney *et al.* (2004) have a North American genotype (Kuehne *et al.*, 2007) that suggests they could have higher optimum growth temperature than *S. paradoxus* with European genotypes (Leducq *et al.*, 2014, 2015). We may therefore underestimate the difference between *S. cerevisiae* and *S. paradoxus* (Leducq *et al.*, 2014). Another laboratory estimate however, suggests that we could be using an overestimate (Salvadó *et al.*, 2011; see Supplemental File 1 for discussion). Thus, the optimum summer temperature range that we predict for *S. cerevisiae* needs to be tested by directly sampling trees in subtropical and tropical regions with precise site locations and trunk girth measurements.

Another important predictor we uncover here for *S. paradoxus* isolation frequency is tree trunk girth (Figure 2), which is consistent with the intuitive notion that older trees harbour a greater diversity of microbial species including yeast. Indeed, the effect of trunk girth is so strong that if we had not included trunk girth in our model, we would not have detected an association of *S. paradoxus* isolation frequency with temperature. Intriguingly, the possible accumulation of yeasts on oak trees as they grow suggests a process of microbial succession that could parallel below ground processes (Bardgett *et al.*, 2005; Bardgett, 2005). Only 42% of the deviance we observed in *S. paradoxus* isolation frequency could be explained by trunk girth and T_max_ together, suggesting that there are other important predictors of *S. paradoxus* isolation frequency that we do not study here. For example, *S. paradoxus* abundance could be influenced by interactions with other microbes (Kowallik *et al.*, 2015); the availability of nutrients (Sampaio and Gonçalves, 2008), water or oxygen (Deak, 2006); by acidity (Deak, 2006) or by sampling season (Glushakova *et al.*, 2007; Charron *et al.*, 2014).

The general caveats that apply when considering climate envelope models (Araújo and Peterson, 2012; Jarnevich *et al.*, 2015) also apply to our findings. We outline regions that have summer temperatures predicted to be associated with high *S. paradoxus* or *S. cerevisiae* isolation frequency (Figure 4). We do not suggest that these regions show the actual distribution of the species however, because they might not contain viable habitat (Araújo and Peterson, 2012; Jarnevich *et al.*, 2015).

Our results also show that *S. paradoxus* and *S. cerevisiae* are not the only oak-associated yeast species with geographic distributions in Europe that could be associated with temperature (Table 1). *W. anomalus* is relevant to humans, as a wine yeast, food spoilage yeast and biocontrol agent (Passoth *et al.*, 2006), occurring naturally on plants, and soil (Kurtzman, 2011). This species can be found on trees in northern North America (Charron *et al.*, 2014; Sylvester *et al.*, 2015) and on central European mountains (Sláviková *et al.*, 2007). We present evidence that *W. anomalus* is more common on northern than on southern European oaks (Table 1), suggesting a southern limit to its distribution in European woodlands. Such a conclusion is consistent with the finding that *W. anomalus* is more often isolated by incubating bark at low than at high temperatures (10°C vs. 30°C; Sylvester *et al.*, 2015). *L. thermotolerans* also naturally occurs on oak bark (Sampaio and Gonçalves, 2008; Charron *et al.*, 2014; Sylvester *et al.*, 2015; Freel *et al.*, 2015) and fruit (Lachance and Kurtzman, 2011), and has been proposed as a good model species for yeast population genetics (Freel *et al.*, 2014, 2015). We find that it is more abundant on oaks in southern Europe (Table 1), consistent with the finding that it is isolated from bark at high temperatures (30°C vs. 10°C; Sylvester *et al.*, 2015).

Knowledge of the climate associations of animal and plant species can lead to the discovery of new populations, as well as the prediction of glacial refugia, biodiversity hotspots, extinction risks and responses to climate change (Araújo and Peterson, 2012; Jarnevich *et al.*, 2015). Because they are too small to see, geographic distributions and therefore ecological associations are more difficult to determine for free-living microbes. However for microbial species that can be cultured, ecologically relevant factors such as temperature preferences are easier to determine experimentally than they are for plants or animals. Our work suggests that laboratory estimates of optimum growth temperature could be used to predict global distributions of free-living microbes.

## Acknowledgements

For field work and field assistance we thank Dan Smith, Cathy Walton, Steve Jones, Ana Maria Pessoa, Carol Smith, Casey Bergman, Will Davenport, Belinda Kemp, Daniela Delneri, Lorenzo Brancia, James Patterson, Richard Harrison, Savvas Vatidis. For contributions in the laboratory we thank Sandra Bunning, Sandra Taylor and Viranga Tilakaratna. For helpful comments on the manuscript we thank Casey Bergman. This work was supported by the Natural Environment Research Council through a NERC Fellowship to DB [grant number NE/D008824/1] (http://www.nerc.ac.uk).

## Authors’ Contributions

D.B. and A.P. conceived and designed the research; A.P. and D.B. performed field sampling; A.P. and H.A.R. performed yeast isolation and species identification; D.B. and H.A.R. analysed the data; D.B. and H.A.R. wrote the manuscript.

## Data Accessibility

DNA sequences determined for this study are available in GenBank: KT206983-KT207282. Photographs of host plants and DNA sequences that did not fulfil the submission criteria at GenBank are available at https://github.com/bensassonlab/yeastecology/.

## Supplemental Files

1. A .pdf with (i) a Supplemental Results section showing the evidence for the isolation of *C. albicans*; (ii) Supplemental Results showing similar *S. cerevisiae* isolation rates from grape, grapevine bark, and oak bark in vineyards, and similar rates from figs and fig tree bark in southern Europe; (iii) Supplemental Results showing that differences among oak trees can explain 52% of the deviance among bark samples, and that bark weight and collection month are not good predictors of the presence of *S. paradoxus*; (iv) Supplemental Results showing the effect of using a different laboratory estimate of the difference in optimal growth temperature for *S. paradoxus* and *S. cerevisiae;* (v) A table of primers used for PCR amplification and DNA sequencing; (vi) Supplemental Figure 1 showing that the approximate geographic positions of *S. cerevisiae* strains from China are close to locations with expected summer temperatures.
2. A .tsv file that summarises the BLAST results for the 371 DNA sequences generated for this study, the species call of the associated yeast strains, and NCBI accession numbers. The query name is the name of the DNA sequence query as it appears in blast outputs; DBuid is the unique identification number in the Bensasson lab yeast collection; ”classification” describes how we classified this sequence for the purpose of our statistical analysis; Ascore is the highest BLAST score when queried against Ascomycota at NCBI; Evalue is the E value associated with this; Cscore and Pscore are the highest BLAST scores when queried against *S. cerevisiae* and *S. paradoxus* respectively. Some DNA sequences were not submitted to NCBI. These were 71 DNA sequences that were technical replicates, contained more than 100 low quality bases (bases with phred-scaled score below 40) or that had fewer than 200 high quality bases and are available at https://github.com/bensassonlab/yeastecology/. Samples with the suffix ”.SM” and ”.YM” for strainUID may contain multiple yeast strains, because they were grown from several colonies each from a Sniegowski selection plate or a YPD plate respectively. All other strains were grown from a single colony.
3. A .tsv file that summarises the presence or absence of *S. cerevisiae* (Scer), *S. paradoxus* (Spar), other yeast that is amplified by primers in the ITS region (otherAmplifiedITS), or other microbial growth (otherGrowth) for every sample collected for this study. This table also includes a description of each sample substrate (e.g. fig, bark, must), field collection date (fieldDate), sample weight (in grams), isolation temperature (°C), the name of the collection site, the species name of the host plant, latitude and longitude (WGS84 format), elevation (in metres), trunk girth (in metres) and pH of soil at base of host where available. Many oak trees classified as most similar to *Q. robur* or *Q. petraea* appeared intermediate between the two species.
4. A .tsv file with details of 301 *S. cerevisiae* and 246 *S. paradoxus* isolates and the geographic locations from which they were sampled. Genotype information is included where it is available. In cases where latitude and longitude were estimated from Google Maps, we include the Google search term used. Where site descriptions cover a large region (e.g. a country name) we selected a point in the centre of that possible region. Yeast isolates with site descriptions that did not allow location within 100-200 km were omitted from this summary. In the case of the *S. paradoxus* strains described in Zhang *et al.* (2010), strain names were not reported, so they are all listed as ”SpNZ”. In Cromie *et al.* (2013), no strains were classified as admixed (or “mosaics”) even though many of the same strains were classified this way in other studies (Liti *et al.*, 2009), we therefore used the data in Cromie *et al.*, 2013 to classify mosaics (those assigned to a single population by InStruct with a probability lower than 0.9375; 15 out of 16 chromosomes). The estimated T*_max_* (in °C×10) for the field site of each strain is shown along with the longitude (TmaxLon) and latitude (TmaxLat) coordinates of the a closest pixel to our estimate of site location at 30 arc-second (approximately 1km) resolution from the WorldClim dataset version 1.4 (1950-2000, release 3, http://www.worldclim.org). However, the positioning of almost all sites is approximate (up to the nearest 100-200km, see Methods).

## References

Almeida, P., Barbosa, R., Zalar, P., Imanishi, Y., Shimizu, K., Turchetti, B., Legras, J.L., Serra, M., Dequin, S., Couloux, A., Guy, J., Bensasson, D., Gonçalves, P., and Sampaio, J.P. (2015). “A population genomics insight into the Mediterranean origins of wine yeast domestication.” Molecular Ecology, page Early View.

Araújo, M.B. and Peterson, A.T. (2012). “Uses and misuses of bioclimatic envelope modeling.” Ecology, 93(7): 1527–1539.

Bardgett, R. (2005). The Biology of Soil: A Community and Ecosystem Approach. OUP Oxford. ISBN 978-0-19-852503-5.

Bardgett, R.D., Bowman, W.D., Kaufmann, R., and Schmidt, S.K. (2005). “A temporal approach to linking aboveground and belowground ecology.” Trends in Ecology & Evolution, 20(11): 634–641.

Bensasson, D. (2011). “Evidence for a high mutation rate at rapidly evolving yeast centromeres.” BMC Evol Biol, 11: 211.

Bensasson, D., Zarowiecki, M., Burt, A., and Koufopanou, V. (2008). “Rapid evolution of yeast centromeres in the absence of drive.” Genetics, 178(4): 2161–2167.

Bonfield, J.K., Smith, K.F., and Staden, R. (1995). “A new DNA sequence assembly program.” Nucleic acids research, 23(24): 4992–4999.

Boynton, PJ., and Greig, D. (2014). “The ecology and evolution of non-domesticated *Saccharomyces* species.” Yeast, 31(12): 449–462.

Charron, G., Leducq, J.B., Bertin, C., Dubé, A.K., and Landry, C.R. (2014). “Exploring the northern limit of the distribution of *Saccharomyces cerevisiae* and *Saccharomyces paradoxus* in North America.” FEMS yeast research, 14(2): 281–288.

Crawley, M.J. (2005). Statistics: An Introduction Using R. Wiley-Blackwell, Chichester, West Sussex, England, 1 edition edition. ISBN 978-0-470-02298-6.

Cromie, G.A., Hyma, K.E., Ludlow, C.L., Garmendia-Torres, C., Gilbert, T.L., May, P., Huang, A.A., Dudley, A.M., and Fay, J.C. (2013). “Genomic sequence diversity and population structure of *Saccharomyces cerevisiae* assessed by RAD-seq.” G3 (Bethesda, Md.), 3(12): 2163–2171.

Deak, T. (2006). “Environmental factors influencing yeasts.” In “Biodiversity and Ecophysiology of Yeasts,” pages 155–174. Springer.

Diezmann, S. and Dietrich, F.S. (2009). “*Saccharomyces cerevisiae:* Population Divergence and Resistance to Oxidative Stress in Clinical, Domesticated and Wild Isolates.” PLoS ONE, 4(4): e5317.

Fay, J.C. and Benavides, J.A. (2005). “Evidence for Domesticated and Wild Populations of *Saccharomyces cerevisiae.*” PLoS Genetics, 1(1): e5.

Fitter, A. and More, D. (2002). Trees. HarperCollins, Glasgow. ISBN 0-00-711074-X 978-0-00-711074-2.

Freel, K.C., Charron, G., Leducq, J.B., Landry, C.R., and Schacherer, J. (2015). “*Lachancea quebecensis* sp. nov., a yeast species consistently isolated from tree bark in the Canadian province of Québec.” International Journal of Systematic and Evolutionary Microbiology.

Freel, K.C., Friedrich, A., Hou, J., and Schacherer, J. (2014). “Population genomic analysis reveals highly conserved mitochondrial genomes in the yeast species *Lachancea thermotolerans*.” Genome Biology and Evolution, 6(10): 2586–2594.

Glushakova, A.M., Ivannikova, Y.V., Naumova, E.S., Chernov, I.Y., and Naumov, G.I. (2007). “Massive isolation and identification of *Saccharomyces paradoxus* yeasts from plant phyllosphere.” Microbiology, 76(2): 205–210.

Goddard, M.R., Anfang, N., Tang, R., Gardner, R.C., and Jun, C. (2010). “A distinct population of *Saccharomyces cerevisiae* in New Zealand: evidence for local dispersal by insects and human-aided global dispersal in oak barrels.” Environmental Microbiology, 12(1): 63–73.

Goddard, M.R. and Greig, D. (2015). “*Saccharomyces cerevisiae:* a nomadic yeast with no niche?” FEMS yeast research, 15(3).

Green, J. and Bohannan, B.J.M. (2006). “Spatial scaling of microbial biodiversity.” Trends in Ecology & Evolution, 21(9): 501–507.

Hijmans, R.J., Cameron, S.E., Parra, J.L., Jones, P.G., and Jarvis, A. (2005). “Very high resolution interpolated climate surfaces for global land areas.” International Journal of Climatology, 25(15): 1965–1978.

Hyma, K.E. and Fay, J.C. (2013). “Mixing of vineyard and oak-tree ecotypes of *Saccharomyces cerevisiae* in North American vineyards.” Molecular Ecology, 22(11): 2917–2930.

Jarnevich, C.S., Stohlgren, T.J., Kumar, S., Morisette, J.T., and Holcombe, T.R. (2015). “Caveats for correlative species distribution modeling.” Ecological Informatics, 29, Part 1: 6–15.

Johnson, L.J., Koufopanou, V., Goddard, M.R., Hetherington, R., Schafer, S.M., and Burt, A. (2004). “Population Genetics of the Wild Yeast *Saccharomyces paradoxus*.” Genetics, 166(1): 43–52.

Kowallik, V., Miller, E., and Greig, D. (2015). “The interaction of *Saccharomyces paradoxus* with its natural competitors on oak bark.” Molecular Ecology, 24(7): 1596–1610.

Kuehne, H.A., Murphy, H.A., Francis, C.A., and Sniegowski, P.D. (2007). “Allopatric divergence, secondary contact, and genetic isolation in wild yeast populations.” Curr Biol, 17(5): 407–11.

Kurtzman, C.P. (2011). “Chapter 80 - *Wickerhamomyces* Kurtzman, Robnett & Basehoar-Powers (2008).” In C.P.K.W.F. Boekhout, editor, “The Yeasts (Fifth Edition),” pages 899–917. Elsevier, London. ISBN 978-0-444-52149-1.

Kurtzman, C.P., Mateo, R.Q., Kolecka, A., Theelen, B., Robert, V., and Boekhout, T. (2015). “Advances in yeast systematics and phylogeny and their use as predictors of biotechnologically important metabolic pathways.” FEMS Yeast Research, 15(6): fov050.

Lachance, M.A., Boekhout, T., Scorzetti, G., Fell, J.W., and Kurtzman, C.P. (2011). “Chapter 90 - *Candida* Berkhout (1923).” In C.P.K.W.F. Boekhout, editor, “The Yeasts (Fifth Edition),” pages 987–1278. Elsevier, London. ISBN 978-0-444-52149-1.

Lachance, M.A. and Kurtzman, C.P. (2011). “Chapter 41 - *Lachancea* Kurtzman (2003).” In C.P.K.W.F. Boekhout, editor, “The Yeasts (Fifth Edition),” pages 511–519. Elsevier, London. ISBN 978-0-444-52149-1.

Leducq, J.B., Charron, G., Samani, P., Dubé, A.K., Sylvester, K., James, B., Almeida, P., Sampaio, J.P., Hittinger, C.T., Bell, G., and Landry, C.R. (2014). “Local climatic adaptation in a widespread microorganism.” Proceedings of the Royal Society B: Biological Sciences, 281(1777): 20132472.

Leducq, J.B., Nielly-Thibault, L., Charron, G., Eberlein, C., Verta, J.P., Samani, P., Sylvester, K., Hittinger, C.T., Bell, G., and Landry, C.R. (2015). “Speciation driven by hybridization and chromosomal plasticity in a wild yeast.” bioRxiv, page 027383.

Liti, G. (2015). “The fascinating and secret wild life of the budding yeast *S. cerevisiae*.” eLife, 4: e05835.

Liti, G., Carter, D.M., Moses, A.M., Warringer, J., Parts, L., James, S.A., Davey, R.P., Roberts, I.N., Burt, A., Koufopanou, V., Tsai, I.J., Bergman, C.M., Bensasson, D., O/’Kelly, M.J.T., van Oudenaarden, A., Barton, D.B.H., Bailes, E., Nguyen, A.N., Jones, M., Quail, M.A., et al. (2009). “Population genomics of domestic and wild yeasts.” Nature, 458(7236): 337–341.

Maganti, H., Bartfai, D., and Xu, J. (2011). “Ecological structuring of yeasts associated with trees around Hamilton, Ontario, Canada.” FEMS yeast research, 12: 9–19.

Martiny, J.B.H., Bohannan, B.J.M., Brown, J.H., Colwell, R.K., Fuhrman, J.A., Green, J.L., Horner-Devine, M.C., Kane, M., Krumins, J.A., Kuske, C.R., Morin, P.J., Naeem, S., Øvreås, L., Reysenbach, A.L., Smith, V.H., and Staley, J.T. (2006). “Microbial biogeography: putting microorganisms on the map.” Nature Reviews Microbiology, 4(2): 102–112.

Naumov, G., Naumova, E., and Sniegowski, P. (1997). “Differentiation of European and Far East Asian populations of *Saccharomyces paradoxus* by allozyme analysis.” Int J Syst Bacteriol, 47(2): 341–344.

Passoth, V., Fredlund, E., Druvefors, U.Ä., and Schnürer, J. (2006). “Biotechnology, physiology and genetics of the yeast *Pichia anomala*.” FEMS Yeast Research, 6(1): 3–13.

Rauch, E.M. and Bar-Yam, Y. (2005). “Estimating the total genetic diversity of a spatial field population from a sample and implications of its dependence on habitat area.” Proceedings of the National Academy of Sciences of the United States of America, 102(28): 9826–9829.

Robert, V., Stalpers, J., Boekhout, T., and Tan, S.h. (2006). “Yeast biodiversity and culture collections.” In “Biodiversity and ecophysiology of yeasts,” pages 31–44. Springer.

Salvadó, Z., Arroyo-López, F.N., Guillamón, J.M., Salazar, G., Querol, A., and Barrio, E. (2011). “Temperature Adaptation Markedly Determines Evolution within the Genus *Saccharomyces*.” Applied and Environmental Microbiology, 77(7): 2292–2302.

Sampaio, J.P. and Gonçalves, P. (2008). “Natural Populations of *Saccharomyces kudriavzevii* in Portugal Are Associated with Oak Bark and Are Sympatric with *S. cerevisiae* and *S. paradoxus*.” Applied and Environmental Microbiology, 74(7): 2144–2152.

Sláviková, E., Vadkertiová, R., and Vránová, D. (2007). “Yeasts colonizing the leaf surfaces.” Journal of Basic Microbiology, 47(4): 344–350.

Sniegowski, P.D., Dombrowski, P.G., and Fingerman, E. (2002). “*Saccharomyces cerevisiae* and *Saccharomyces paradoxus* coexist in a natural woodland site in North America and display different levels of reproductive isolation from European conspecifics.” FEMS Yeast Research, 1(4): 299–306.

Sutton, D.A. (1990). Trees of Britain and Europe. Kingfisher, London. ISBN 0-86272-523-2 978-0-86272-523-5.

Sweeney, J.Y., Kuehne, H.A., and Sniegowski, P.D. (2004). “Sympatric natural *Saccharomyces cerevisiae* and *S. paradoxus* populations have different thermal growth profiles.” FEMS Yeast Research, 4(4-5): 521–525.

Sylvester, K., Wang, Q.M., James, B., Mendez, R., Hulfachor, A.B., and Hittinger, C.T. (2015). “Temperature and host preferences drive the diversification of Saccharomyces and other yeasts: a survey and the discovery of eight new yeast species.” FEMS yeast research, 15(3): fov002.

Tanghe, A., Carbrey, J.M., Agre, P., Thevelein, J.M., and Van Dijck, P. (2005). “Aquaporin expression and freeze tolerance in *Candida albicans*.” Applied and Environmental Microbiology, 71(10): 6434–6437.

Taylor, J.W., Turner, E., Townsend, J.P., Dettman, J.R., and Jacobson, D. (2006). “Eukaryotic microbes, species recognition and the geographic limits of species: examples from the kingdom Fungi.” Philosophical Transactions of the Royal Society B: Biological Sciences, 361(1475): 1947–1963.

Wang, Q.M., Liu, W.Q., Liti, G., Wang, S.A., and Bai, F.Y. (2012). “Surprisingly diverged populations of *Saccharomyces cerevisiae* in natural environments remote from human activity.” Molecular Ecology, 21(22): 5404–5417.

Will, J.L., Kim, H.S., Clarke, J., Painter, J.C., Fay, J.C., and Gasch, A.P. (2010). “Incipient Balancing Selection through Adaptive Loss of Aquaporins in Natural *Saccharomyces cerevisiae* Populations.” PLoS Genet, 6(4): e1000893.

Zhang, H., Skelton, A., Gardner, R.C., and Goddard, M.R. (2010). “*Saccharomyces paradoxus* and *Sac-charomyces cerevisiae* reside on oak trees in New Zealand: evidence for migration from Europe and interspecies hybrids.” FEMS Yeast Research, 10(7): 941–947.Tables and Figures

